# STING activation promotes robust immune response and NK cell-mediated tumor regression in glioblastoma models

**DOI:** 10.1101/2022.02.28.481908

**Authors:** Gilles Berger, Erik H. Knelson, Jorge L. Jimenez-Macias, Michal O. Nowicki, Saemi Han, Eleni Panagioti, Patrick H. Lizotte, Kwasi Adu-Berchie, Alexander Stafford, Nikolaos Dimitrakakis, Lanlan Zhou, E. Antonio Chiocca, David J. Mooney, David A. Barbie, Sean E. Lawler

## Abstract

Immunotherapy has had a tremendous impact on cancer treatment in the past decade, with hitherto unseen responses at advanced and metastatic stages of the disease. However, the aggressive brain tumor glioblastoma (GBM) is highly immunosuppressive and remains largely refractory to current immunotherapeutic approaches. The cGAS-STING cytoplasmic double stranded DNA (dsDNA) sensing pathway has emerged as a next-generation immunotherapy target with potent local immune stimulatory properties.

Here, we investigated the status of the STING pathway in GBM and the modulation of the brain tumor microenvironment (TME) with the STING agonist ADU-S100. Our data reveal the presence of STING in human GBM specimens, where it stains strongly in the tumor vasculature. We show that human GBM explants can respond to STING agonist treatment by secretion of inflammatory cytokines. In murine GBM models, we show a profound shift in the tumor immune landscape after STING agonist treatment, with massive infiltration of the tumor-bearing hemisphere with innate immune cells including inflammatory macrophages, neutrophils and NK populations. Treatment of established murine intracranial GL261 and CT-2A tumors by biodegradable ADU-S100-loaded intracranial implants demonstrated a significant increase in survival in both models and long-term survival with immune memory in GL261. Responses to treatment were abolished by NK cell depletion. This study reveals therapeutic potential and deep remodeling of the TME by STING activation in GBM and warrants the further examination of STING agonists alone or in combination with other immunotherapies such as cancer vaccines, CAR T cells, NK therapies or immune checkpoint blockade.

**Significance statement:** Modulation of the immune microenvironment is critical for immunosuppressive and therapy refractory tumors like glioblastoma. Activation of the STING pathway deeply remodels the brain tumor environment and attracts innate immune cells and natural killer cell populations, producing a robust antitumor effect with long-term immune memory. We further show that human glioblastoma tissue can respond to the therapy and lay the foundations for combined intracranial immunotherapies by using crosslinked biodegradable brain implants.

## Introduction

Immunotherapy has profoundly altered cancer treatment (1, 2). In particular, unprecedented responses to immune checkpoint blockade (ICB) in some cancer types have clearly established that the host immune system can be retrained to eliminate tumors. However, many tumors remain resistant to ICB and numerous studies suggest that these may benefit from additional treatments that create a tumor microenvironment more conducive to immune activation (3, 4). Therefore, understanding the key mechanisms needed to effectively modulate the intratumoral microenvironment is an area of major importance, and therapeutics that break local intratumoral immunosuppressive mechanisms may allow the development of effective anti-tumor immunity.

Glioblastoma (GBM) is the most common primary malignant brain tumor, with approximately 10,000 newly diagnosed cases per year in the U.S. (5). Patients have a dismal median survival of 15 months with the current standard of care of surgery followed by post-operative chemo-radiotherapy (6), and new therapeutic approaches are still of an urgent and unmet need. Despite the successes of ICB in some cancers (7–9), GBM remains resistant, albeit with some indications of response in the neoadjuvant setting in recurrent GBM (10–14). It is thought that highly immunosuppressed ‘cold’ and non-immunogenic tumor microenvironment (TME) in GBM is a major factor in resistance to ICB (13). Local immunostimulatory approaches can enhance ICB efficacy in GBM in preclinical settings (15, 16), and clinically in other tumor types (17–19). These employ the intratumoral (IT) delivery of agents like oncolytic viruses (20, 21) or small molecules that activate innate immune signaling in the TME, with the goal of initiating an anti-tumor immune response, overcoming immunosuppressive mechanisms and remodeling the TME (22, 23).

The cGAS-STING cytoplasmic double-stranded DNA (dsDNA) sensing pathway has emerged as a next-generation immunotherapy target with potent local immune stimulatory properties. The stimulator of interferon genes (STING) protein is localized to the endoplasmic reticulum membrane and is critical for immune-sensing of pathogens and cancer. Activation of STING leads to type-I interferon (IFN) production in response to cytosolic dsDNA (24–27). The sensor protein for cytosolic dsDNA is the enzyme cGAS, which catalyzes the formation of the cyclic dinucleotide (CDN) cyclic-GMP-AMP ([G(2′,5′)pA(3′,5′)p]; cGAMP) (28–34). These CDNs bind the STING dimer (32), inducing conformational changes and downstream events leading to the recruitment and phosphorylation of TANK binding kinase 1 (TBK1) followed by the dimerization and phosphorylation of interferon regulatory factor 3 (IRF-3) and the transcription of interferon-associated genes (31, 35–38). Besides the endogenous 2’3’-cGAMP, CDNs can be pathogen-derived and are ubiquitous second messengers in prokaryotic species (23).

ADU-S100, a synthetic compound used in the present study, is based on the typical CDN scaffold, with two adenines and a substitution of the phosphodiester linkages by phosphorothioates, making it resistant to enzymatic degradation (39, 40). STING agonists promote potent anti-tumor immunity in preclinical models (25, 26, 41–43), are considered promising anticancer agents with remarkable preclinical efficacy in some tumor models (21, 22, 24, 44, 45), and are being investigated in clinical trials for various solid cancers. STING activation in the brain for cancer treatment has also shown promise in initial studies (46–48), but the nature of the STING pathway in tumors like GBM has not been delineated and the effects of STING agonists on the GBM TME have not been explored in detail. Their application in the central nervous system (CNS) and for GBM treatment are thus still poorly defined but have potential to overcome the high levels of immunosuppression in GBM (46).

Here, we show that STING can be activated in human GBM, where it is expressed highly in tumor-associated blood vessels. We define responses to intratumoral (IT) STING agonist delivery in murine GBM models, and show that IT biodegradable implants loaded with ADU-S100 can promote long-term survival and immune memory in murine GBM, supporting further development of this approach.

## Results

### STING pathway status in GBM

Although STING is considered a promising target in cancer, expression of STING pathway components in GBM has not been studied. Therefore, we first characterized the expression of the key components of the cGAS-STING-IRF3 pathway in GBM, both in patient GBM samples and GBM cell lines. Western blotting showed low levels of cGAS expression in patient GBM samples and various levels of STING, TBK1 and IRF3 (Fig 1a). All the GBM cell lines tested, comprising patient-derived neurosphere cells G9 and G30 (49), as well as the two murine models used in the present work (GL261 and CT-2A), express cGAS, STING, TBK1 and IRF3 at the protein level (Fig. 1a). Analysis of STING and phospho-TBK1 (Ser172) immunohistochemical staining in a tissue microarray (TMA) showed a range of expression which is highest in GBM and lowest in normal brain with intermediate levels in anaplastic astrocytoma and oligodendroglioma specimens (Fig. 1b). In the normal human brain, STING is expressed in a subset of cells and is particularly prominent in the vasculature, and this picture holds true for the majority of GBM cases in our TMA (Fig. 1c and d). Staining was also observed in individual cells scattered throughout the tumor parenchyma. pTBK1 was detected in the vasculature of GBM samples, indicating that there may be some STING pathway activation in GBM vasculature in contrast to normal brain vasculature where pTBK1 staining is absent (Fig. 1c,d) (50). In mice, STING is readily detectable in both CT-2A and GL261 tumors *in vivo* (Supplementary Figure S1). The cGAS-STING pathway shows some degree of baseline activation in GL261 tumors, pTBK1 being colocalized with markers of the vasculature such as CD31 and α-SMA (Fig. 1e). Tumors are also infiltrated and surrounded by STING positive cells (Fig. 1f and g) that mainly comprise F4/80^+^ and IBA1^+^ cells, as members of the innate myeloid immune population and microglia (Fig. 1h and i, respectively).

**Fig. 1.**
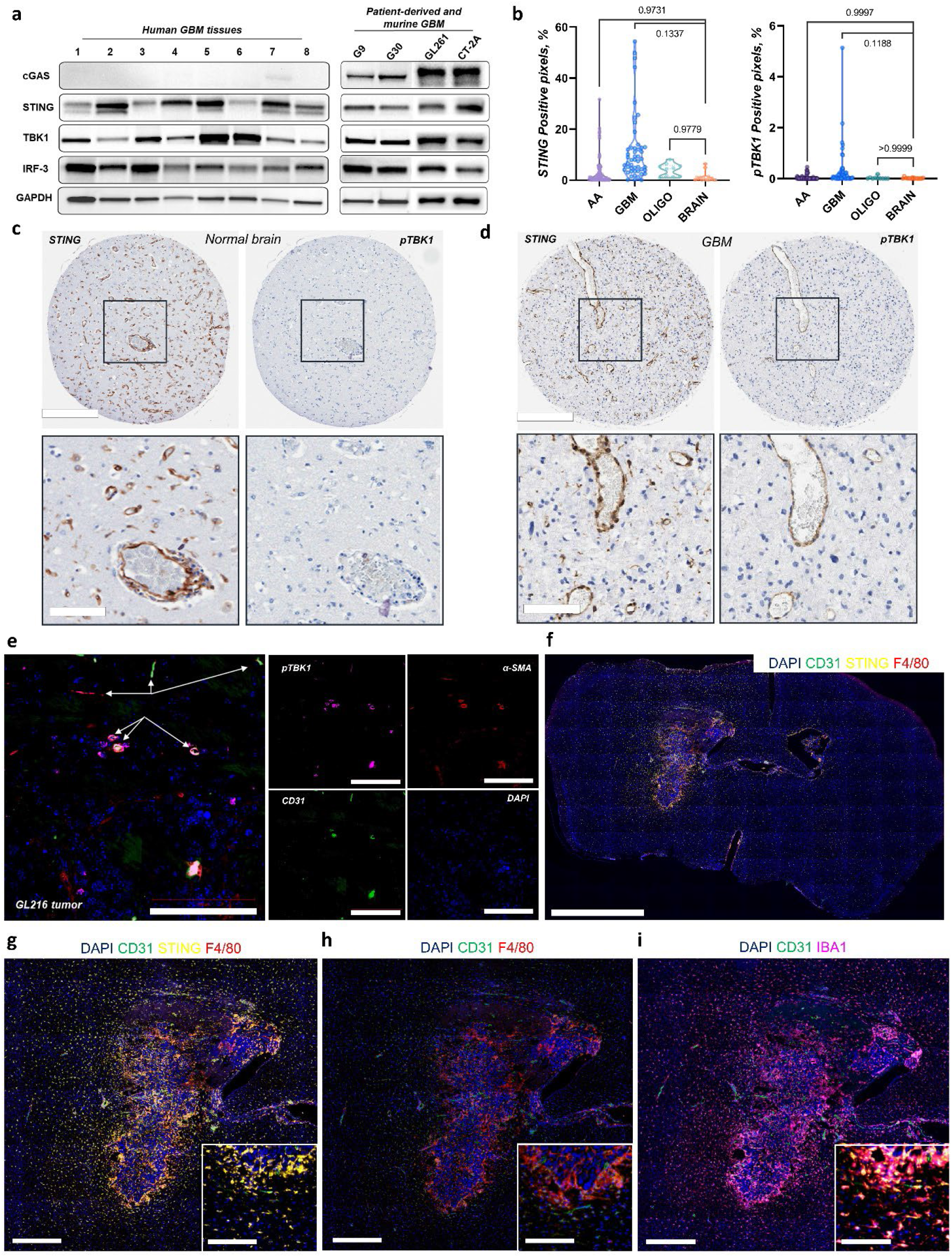
Assessment of STING pathway expression in glioblastoma. **a**, Immunoblot of cGAS, STING, TBK1, IRF-3 and GAPDH levels in patient GBM tissue, patient-derived GBM cells and murine GL261 and CT-2A glioma cells. **b**, Quantification of tissue microarray (TMA) sections for STING and pTBK1, as percentage of positive pixels. AA, n = 77; GBM, n = 49; oligo, n = 10; brain, n = 16. *P* values calculated by one-way ANOVA. **c**, Representative TMA sections immunostained for STING and pTBK1 in normal brain and **d**, GBM. Scale bars = 400 and 100 µm. **e**, Immunofluorescence staining showing partial activation of the STING cascade in GL261 tumors and the colocalization of pTBK1 within the vasculature (CD31 and α-SMA). Left panel: white arrows point to blood vessels. Right panel: Split channels show colocalization of pTBK1 with the vascular markers CD31 and α -SMA. Scale bar = 100 µm. **f**, Multiplex immunofluorescence staining on a whole brain section from a GL261 tumor bearing mouse showing DAPI (blue), CD31 (green), STING (yellow) and F4/80 (red). Scale bar = 1 mm. **g, h** and **i**, Immunofluorescence staining of the tumor zone showing selected markers as indicated. Scale bars = 400 and 100 µm.

### The STING pathway is functional in GBM and elicits immune-mediated tumor cell killing in vitro

CXCL10 is an important cytokine produced downstream of type I IFN after STING activation and is commonly used to measure STING activity in human cancer models (51, 52). Using a CXCL10 ELISA, we found that the STING pathway is non-functional in all of the tested human GBM cell lines (Supplementary Figure S2). Similarly, the murine glioma cell lines CT-2A and mut3 were not responsive to STING agonists, with GL261 cells being a notable outlier which responded strongly to STING agonist treatment as demonstrated by CXCL10 release (Fig. 2a). Human brain vascular pericytes (HBVP) and the human brain microvascular endothelial cell line hCMEC/D3 were responsive to STING agonist treatment supporting a role of STING in the vasculature (Fig. 2a). We then investigated the feasibility of activating STING in GBM, using patient GBM specimens that were cultured as explants in suspension, and CXCL10 release measured after treatment with the STING agonist ADU-S100 (50). These patient samples were responsive to treatment with the STING agonist, producing various levels of inflammatory cytokines, indicating the activation of the STING pathway within the tumor tissue (Fig. 2b and c), and establishing that STING can be activated in human GBM and is therefore a potential immunotherapeutic target. After having established that the STING pathway was present and functional in GBM tissue, we tested the activity of STING agonists *in vitro* through co-culture immune-mediated cell killing assays (Fig. 2d and e). GBM neurospheres made from GFP-expressing G9pCDH patient-derived GBM cells (49) were incubated with human PBMCs freshly extracted from healthy donors at different ratios and treated with ADU-S100 (Fig. 2e). This showed that ADU-S100 is not toxic to GBM neurospheres, even at the highest concentration (Fig. 2e). In contrast, in the presence of PBMCs we observed an ADU-S100 concentration-dependent immune-cell killing of the GBM neurospheres (Fig. 2f) with immune-mediated cytotoxicity being efficient between 12.5 and 50 µM, while the effect is reduced at 100 µM probably due to T cell toxicity at high STING agonist concentrations, as evidenced previously (53). The effects of the STING agonist increased with PBMC concentration indicating an immune-mediated killing effect (Fig. 2f).

**Fig. 2.**
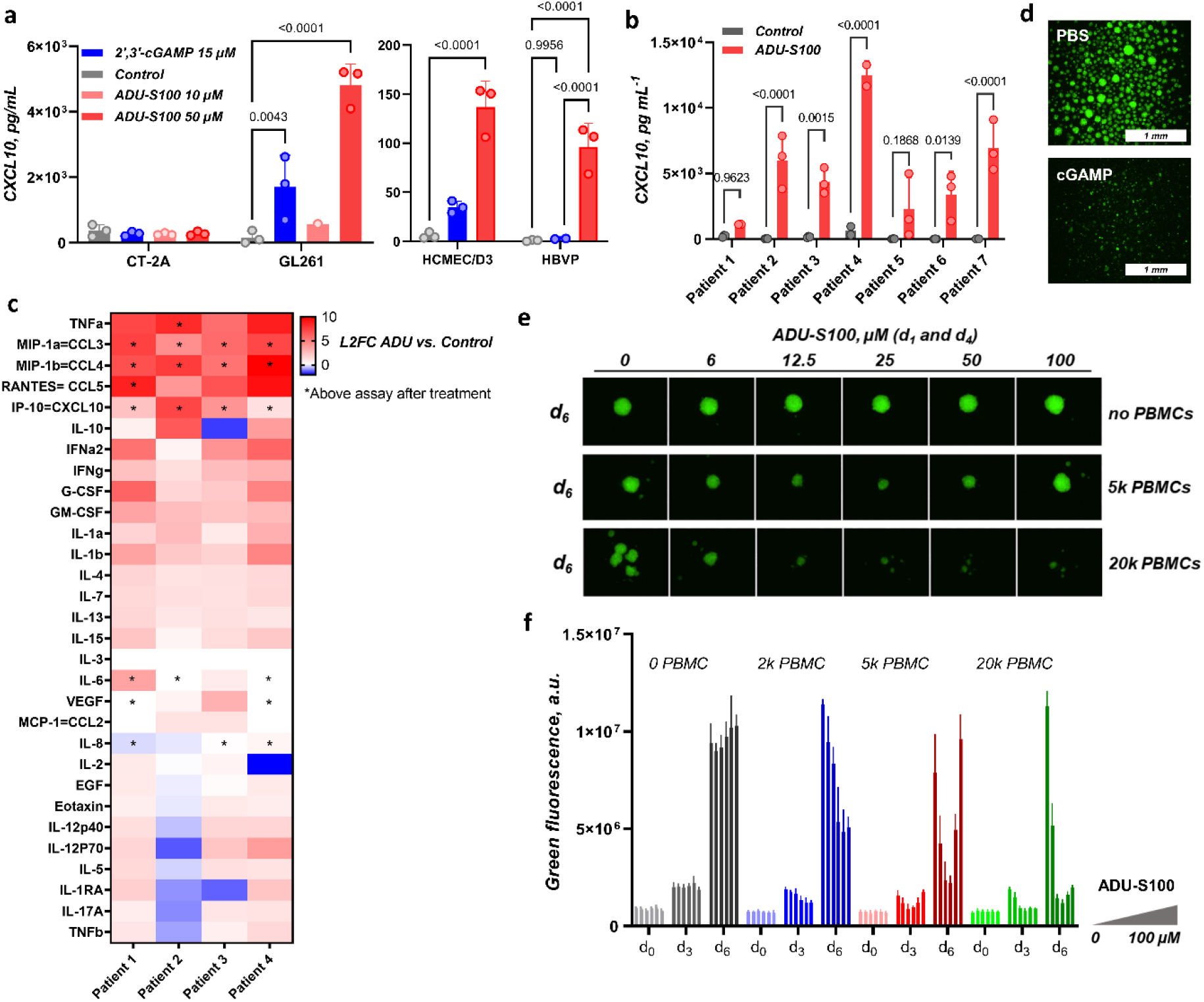
Effects of STING activation on GBM cells. **a**, Levels of CXCL10 as measured by ELISA 24 h after STING agonist treatment of the indicated cell lines. *P* values calculated by two-way ANOVA. **b**, CXCL10 ELISA levels from freshly resected patient GBM specimens cultured for 2-5 days post-surgery (patient 1: 2 days, patients 2, 3, 5, 6 and 7: 3 days, patient 4: 5 days). Control vs ADU-S100 (50 µM). *P* values calculated by two-way ANOVA. **c**, Log_2_ fold-change cytokine/chemokine differences in conditioned media ADU-S100 (50 µM) vs controls, from freshly resected patient GBM specimens cultured as above. **d**, PBMC/G9pCDH co-culture, ± cGAMP treatment at 100 µM, formulated with lipofectamine. **e**, Immuno-GILA assay by co-culture of fluorescent neurospheres (G9pCDH) and fresh PBMCs, treated with ADU-S100 (0 to 100 µM). **f**, Fluorescence plots from the immuno-GILA assay with the various GSC:PBMC ratios at the indicated cell numbers over a period of days as shown in the bar chart over a range of drug concentrations.

### STING activation in intracranial GBM models drives innate immune cell infiltration

To understand the effects of STING agonists on the GBM TME, we characterized the immune response and immune infiltrates after ADU-S100 treatment of GL261 and CT-2A tumors in immunocompetent mice. Initial pilot studies using 2’,3’-cGAMP or ADU-S100 to activate STING indicated a strong innate immune response three days after treatment and an increase in myeloid populations, together with a survival benefit (Supplementary Figure S3). We then analyzed in detail brain infiltrating leukocytes (BILs) by flow cytometry using a panel of 13 immunological markers after an intratumoral (IT) bolus of 50 µg ADU-S100 (39, 54) at day 14 post GL261 intracranial implantation. Three days after treatment, we observed the suppression of microglia, and both CD4+ and CD8+ T cell populations, and infiltration of NK and CD11b^+^/Gr1^+^ inflammatory immune cells (Fig. 3b). At 7 days post-treatment T cell proportions recovered and were increased compared with baseline levels. PD-L1 expression was sharply increased shortly after treatment in CD45-negative cells (Fig. 3c, day 3). In this regard, G-MDSC populations also increase, although these may not be mature and immunosuppressive, but rather inflammatory (Fig. 3d). These populations do not show increased expression of PD-L1 (Supplementary Fig. S4a).

**Fig. 3.**
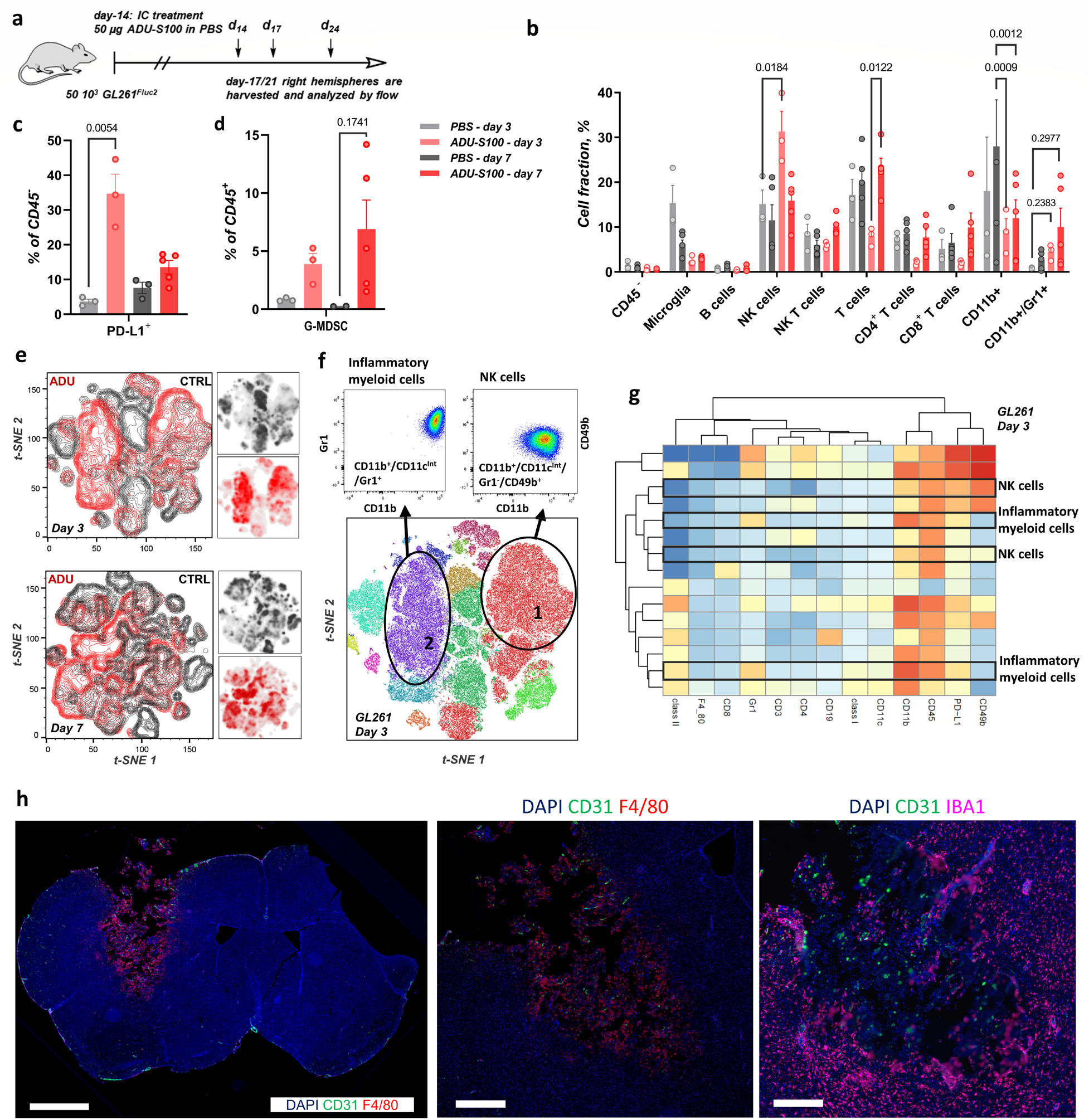
Assessment of GL261 tumor immune infiltrates after STING agonist treatment. **a**, Timeline of the GL261 *in vivo* experiments for BILs. Biologically independent animals per group (ADU-S100/PBS day 3, n = 3; PBS day 7, n = 3; ADU-S100 day 7, n = 5). *P* values calculated by two-way ANOVA. **b**, BIL profile of GL261 tumors at days 3 and 7 using a typical gating procedure. **c**, Percentage of PD-L1^+^ CD45^-^ cells. *P* value calculated by unpaired t-test. **d**, G-MDSC populations within the BILs. *P* value calculated by unpaired t-test. **e**, 2D t-SNE plots at day 3 and 7, treated mice in red and controls in dark grey. **f**, t-SNE map for treated mice at day 3 colored by the FlowSOM populations. **g**, Heatmap and hierarchical clustering of the FlowSOM populations at day 3. **h**, Immunofluorescence staining on a whole brain section and the necrotic tumor zone 72 h after ADU-S100 treatment (50 µg, bolus in PBS).

To obtain both a more global and precise picture of immune activation following IT ADU-S100 treatment in the brain, we analyzed the multidimensional data by t-SNE coupled to FlowSOM (55), allowing efficient clustering of populations and visualization of the global shift in immune populations for treated samples. Figure 3e shows 2D t-SNE density plots of the controls and ADU-S100-treated groups at day 3; the immunological landscape reveals extensive reorganization of the TME after treatment and activation of the STING pathway (Fig. 3e). The TME at day 7 also follows a deep remodeling (Fig. 3e). We proceeded to cluster these populations using FlowSOM and this revealed that the cell types making the bulk of the immune profile of the treated brain infiltrates were of two main types, which are mapped on the tSNE plot at day 3 in Figure 3f: (1) NK populations, defined as CD3^-^/CD11b^+^/CD11c^Int^/Gr1^-^/CD49b^+^ and (2), inflammatory myeloid cells comprising macrophages (MΦ), dendritic cells (DCs) and neutrophils (N), collectively defined as CD3^-^/CD11b^+^/CD11c^Int^/Gr1^+^/CD49b^-^. These could also comprise MDSC-type populations, although in this case represent immature myeloid cells that would not yet convey suppressive phenotypes (56). These inflammatory populations are seen in other inflammatory states in the brain, such as in the response to traumatic brain injury (57). Figure 3g shows the clustering of the FlowSOM populations at day 3 for GL261, and the main increased cell populations are highlighted. We can divide these into two major groups, one belonging to the NK type, with low Gr1 and mid-to high expression of CD49b, and the other group comprising Gr1^+^/CD49b^-^ inflammatory populations. Cytotoxic infiltrates of F4/80^+^ cells are further evidenced by the multiplexed IF images 72 h after treatment of GL261 tumors (Fig. 3h).

We then performed a similar study in CT-2A tumors treated with a 50 µg ADU-S100 bolus in PBS (day 7 post-implantation) and the tumor-bearing hemispheres were analyzed for BILS (Fig. 4a). As in the case of GL261, we observed upregulation of PD-L1 and changes in the MDSC populations, the latter however do not show significant changes in PD-L1 expression (Fig. 4b and Supplementary Figure S4b). 2D t-SNE once again revealed a complete remodeling of the TME and its immune profile, with density plots that are completely shifted (Fig. 4c); showing a consistent and significant activation of the innate immune system at day 3, with an inflammatory TME comprised of NK populations and inflammatory immature innate cells of the myeloid lineage (Fig. 4d, e and f). A similar picture is obtained at day 7 with NK and innate immune cell infiltrates (Supplementary Figure S5). Our results in the brain therefore support the critical participation of NK populations in the STING-induced antitumor effects (58, 59). Thus, intratumoral delivery of STING agonists in murine GBM results in a strong innate immune response characterized by partially immature myeloid infiltrates and PD-L1 upregulation.

**Fig. 4.**
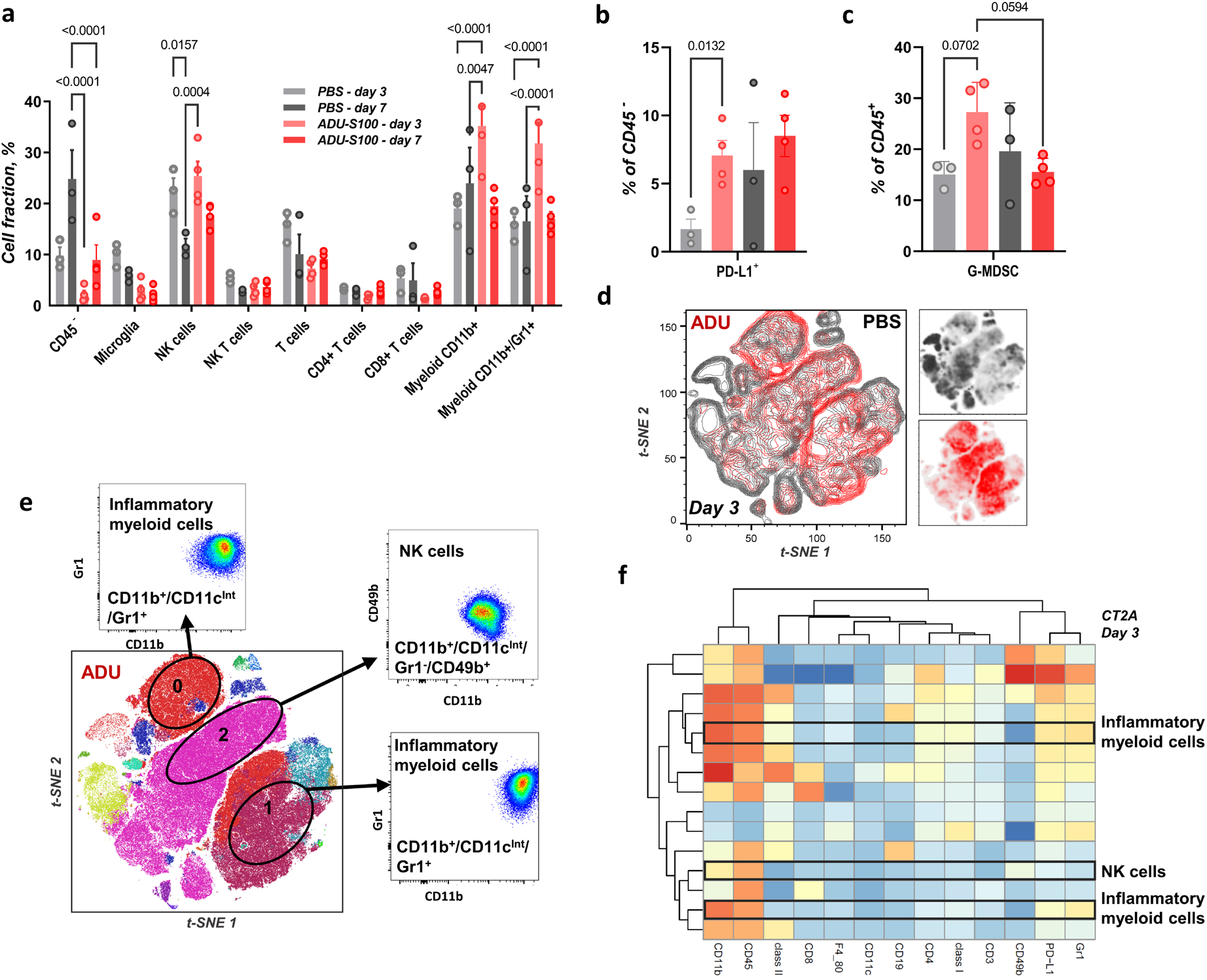
Assessment of CT-2A tumor immune infiltrates after STING agonist treatment. **a**, BILs profile of CT-2A tumors at days 3 and 7 using a typical gating procedure. Biologically independent animals per group (PBS, n = 3; ADU-S100, n = 4). *P* values calculated by two-way ANOVA. **b**, PD-L1^+^ percentage of CD45^-^ cells. *P* value calculated by unpaired t-test. **c**, MDSC populations within the BILs. *P* values calculated by one-way ANOVA. **d**, 2D t-SNE plots at day 3, treated mice in red and controls in dark grey. **e**, t-SNE map for treated mice at day 3 colored by the FlowSOM populations; main upregulated FlowSOM populations are highlighted. **f**, Heatmap and hierarchical clustering of the FlowSOM populations at day 3.

We proceeded to analyze the transcriptome of GL261 tumors after ADU-S100 treatment and isolation of the BILs from the tumor-bearing hemisphere at 72 h. We can clearly distinguish the two conditions and the effect of the STING therapy, as seen from the principal component analysis (PCA) on the individual samples (Fig. 5a). This showed an acute specific activation of innate defense mechanisms (Fig. 5b and d). A volcano plot highlighted 20 of the most differentially expressed genes, showing high induction of the *Ifit1* gene (interferon-induced protein with tetratricopeptide repeats 1B-like 1), in accordance with the activation of the type I IFN pathway. IFIT proteins are important for viral defense and are known to be widely expressed in the mouse brain. Activation of the *ifit1* gene downstream of DNA sensing has been established (60). Other highly overexpressed genes included *Ifi47, Tnf* and *Mx2*, all of which have been implicated in IFN signaling (Fig. 5b) (61). It is worth noting that the TNFα response is also seen from our human Luminex panel on the fresh GBM samples (Fig. 2c). Gene enrichment analysis also revealed terms related to the type I IFN signaling pathway being highly enriched in the ADU-S100 treated samples, in addition to terms related to NF-kB activation and translocation, cytokine production and inflammatory response, while the lipoxygenase pathway was downregulated (Fig. 5d) (62), further supporting immune activation. We also compared gene expression signatures between ADU-S100 treated and PBS controls for genes known to be responsive to IFNγ (type II IFN), IFNβ (type 1 IFN) and IFNα (type 1 IFN) treatment (Fig. 5c) (63). Treated samples had higher average gene expression signatures for all three IFN responsive gene sets that were investigated, with IFNα and IFNβ showing statistically significant differences between the two conditions (FDR adjusted p values of 0.0062 and 0.0032, respectively). Transcriptomic analyses therefore showed that the GL261 tumors were highly responsive to ADU-S100 treatment, characterized by potent IFN signaling and innate immune activation. The involvement of NK cells in the response to the STING agonist was highlighted by increased NK cell gene signature shown in the violin plot (Fig. 5c). NCR1, KLRD1, CD247, PRF1 and TNF are found among the highly upregulated genes after ADU-S100 treatment. NCR1 (NKp46) is the most highly upregulated and is one of the main activators of NK cells (64, 65). KLRD1, also known as CD94, is widely expressed in NK cells and some T cells, and functions together with NKG2A/C as an immune checkpoint (66). CD247, expressed in NK and T cells, associates with NCR1, NCR3 and CD16. It is involved in the responsiveness of NK cells, e.g. degranulation (66). PRF1 (perforin 1), is a major component of cytolytic granules and is involved in the cytotoxic activity of NK and T cells (66). A volcano plot for this specific gene set (*i*.*e*., NK-mediated cytotoxicity) can be found in the Supplementary Figure S6.

**Fig. 5.**
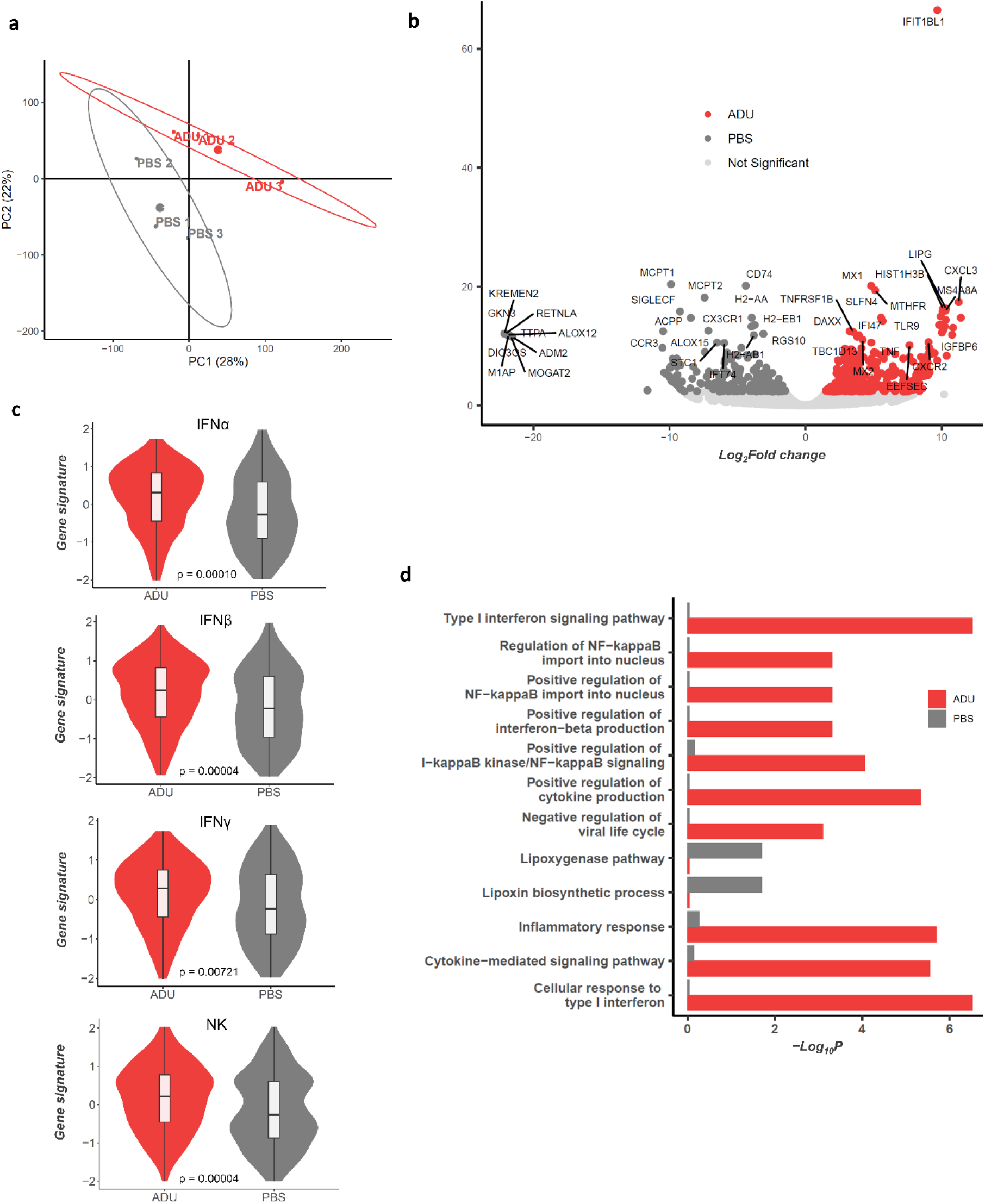
Transcriptome analysis from GL261 tumors treated with ADU-S100. **a**, PCA plot for individual samples, ellipsoids drawn with their centroids at the 95% confidence interval. The first two PCs are shown with their proportion of variance explained. Biologically independent animals per group, n = 3. **b**, Volcano plots for RNA sequencing data from GL261 BILs, ADU-S100 vs PBS showing differentially expressed genes (FDR adjusted *p* threshold ≤ 0.1.) **c**, Violin plots showing aggregate expression of IFN and NK-mediated cytotoxicity genes after ADU-S100 treatment of GL261 tumors. FDR adjusted *P* values (Wilcoxon) are given for each gene set (ADU-S100 vs PBS). **d**, Gene ontology analysis after ADU-S100 treatment of GL261 tumors.

### Therapeutic STING brain implants show high efficacy in mouse models

With evidence of immune-dependent tumor cell killing, remodeling of the TME and elevated innate immune response after IT injection of ADU-S100, we performed efficacy studies in murine GBM models. This was performed using ADU-S100 loaded intratumoral implants, made of cross-linked alginate chains and designed to release the STING agonist over a period of a few days (Fig. 6a and b) (67). This approach was chosen as it may be more clinically applicable than the bolus injection and can also be adapted and combined with other therapies such as immune checkpoint blockade. The gel matrix used here has been composed to perform as a rapid release system for small molecules, and a slow release delivery for larger molecules. As such, the small molecule STING agonist is released quickly, to mimic the bolus injection and trigger the immune reaction, while the aPD-1 antibody will slowly come to effect to oppose the chronic immunosuppressive effects of STING activation over time. To test the hydrogel approach, GL261 tumors were treated with 50 µg ADU-S100 in biodegradable hydrogels two weeks post-implantation and the tumors were monitored by bioluminescence and magnetic resonance imaging (MRI) starting 4 days after treatment (Fig. 6c, d and e). A striking effect of the therapy was seen shortly after treatment with the IVIS signal increasing sharply in the control group (Fig. 6e), together with the tumor size as seen from MRI (Fig. 6d, for full MRI data, see Supplementary Figure S7). The tumors completely regressed two weeks after treatment in two-thirds of the treatment group while the mock treated mice were already beyond their survival endpoints. The observed survival rate of STING monotherapy is thus comparable to the reported combined aPD-1 and aCTLA-4 therapy in the same model (18). Two-thirds of the treated mice were long-term survivors and were rechallenged by injection of GL261 cells into the contralateral hemisphere at day 150 and did not develop tumors, showing the establishment of long-term adaptive immunity after STING agonist treatment of GL261 tumor-bearing animals (Fig. 6c). Analysis of BILs 17 days after treatment with our ADU-S100 implants, showed a sustained myeloid infiltration and a low PD-L1 expression on CD45-negative cells, in contrast with the acute upregulation shortly after treatment (Supplementary Figure S8c and d). The treated mice were tumor-free at that time point and went on to become long-term survivors.

**Fig. 6.**
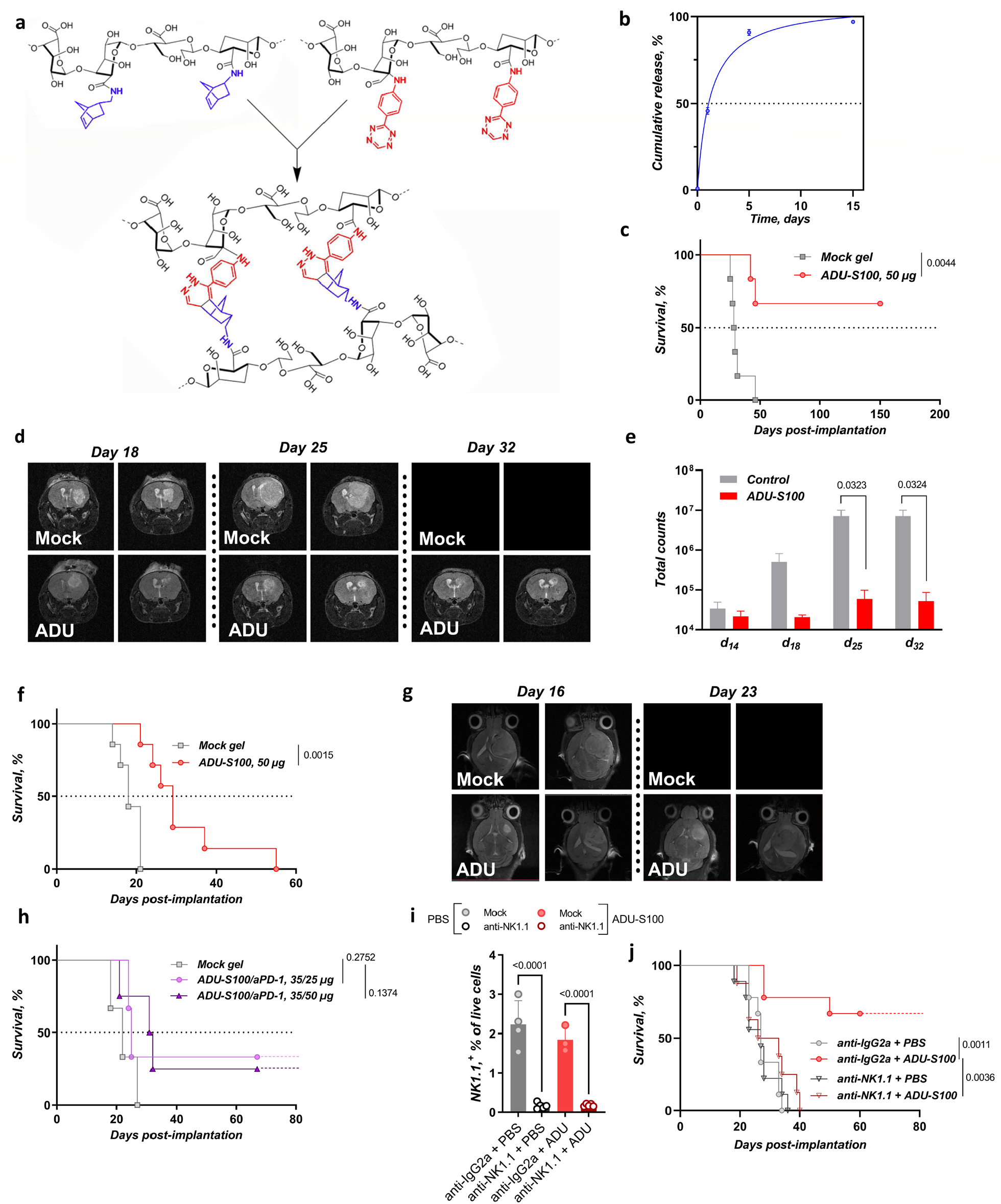
STING agonist treatment of murine GBM *in vivo*. **a**, Biodegradable cross-linked hydrogels used as intracranial implants. **b**, *In vitro* gel release profile for cGAMP. **c**, Kaplan-Meier plot for the GL261 ADU-S100 monotherapy study. *P* value calculated by the Gehan-Breslow-Wilcoxon test. 5×10^4^ GL261^Fluc^ cells were implanted at day 0 and intracranial gels were implanted at day 14. Biologically independent animals per group (mock gel, n = 6; ADU-S100 50 µg, n = 6). **d**, GL261 *in vivo* efficacy study showing MRI starting day 18 post-treatment. Blank images represent euthanized animals. **e**, Bioluminescence IVIS signal from treatment day to endpoint, GL261^FLuc^. *P* values calculated by multiple unpaired t-tests. **f**, Kaplan-Meier plot for the CT-2A ADU-S100 monotherapy study. *P* value calculated by the Gehan-Breslow-Wilcoxon test. 5×10^4^ CT-2A cells were implanted at day 0 and intracranial gels were implanted at day 7. Biologically independent animals per group (mock gel, n = 6; ADU-S100 50 µg, n = 7). **g**, CT-2A *in vivo* efficacy study showing representative MRI pictures at days 18 and 23. Blank images represent euthanized animals. **h**, Kaplan-Meier plot for the CT-2A ADU-S100/aPD-1 combination study. *P* value calculated by the Gehan-Breslow-Wilcoxon test. 5×10^4^ CT-2A cells were implanted at day 0 and intracranial gels were implanted at day 7. Biologically independent animals per group (mock gel, n = 3; ADU-S100/aPD-1 35/25 µg, n = 3; ADU-S100/aPD-1 35/50 µg, n = 4). **i**, Percentage of NK cells as measured in the whole blood of mice 32 h after intraperitoneal administration of the anti-NK1.1 depleting antibody, leading to >98 % removal of the targeted immune cells (n = 3-5 mice per group). *P* values calculated by one-way ANOVA. **j**, Kaplan-Meier plot for the GL261 ADU-S100 monotherapy study with and without NK depletion. 5×10^4^ GL261^Fluc^ cells were implanted at day 0 and mice were treated intracranially at day 14. The depleting antibody was injected 36 h before STING agonist treatment and biweekly after that. *P* value calculated by the Gehan-Breslow-Wilcoxon test. Biologically independent animals per group (mock + PBS, n = 8; anti-NK1.1 + PBS, n = 8; mock + ADU-S100 50 µg, n = 9; anti-NK1.1 + ADU-S100 50 µg, n = 8).

We then turned to the aggressive and notably colder, non-immunogenic CT-2A model as a more challenging syngeneic GBM mouse model (68, 69). The high immunogenicity of GL261 tumors may not correlate well with the clinical reality of GBM (70, 71). A similar efficacy study using cross-linked hydrogels to deliver ADU-S100 was conducted with the CT-2A model. This showed a significant increase in the median survival from 18 to 29 days (Fig. 6f). Representative MRI pictures are shown in Fig 6g and the full set of images can be found in Supplementary Figure S9. No long-term survivors were observed using the STING monotherapy, supporting previous observations that the CT-2A model is more resistant to immunotherapy than GL261 (68). We then performed an initial pilot study of the combination of ADU-S100 and aPD-1 loaded into hydrogels. This led to the emergence of long-term survivors (Fig. 6h), supporting further detailed studies with this approach.

Finally, to evidence the critical involvement of NK populations in the tumor cell killing and the preclinical efficacy of ADU-S100 treatment in our models, we performed a NK depletion study on GL261-bearing mice. NK cell depletion was performed prior to treatment (Fig. 6i) and throughout follow-up, showing a complete loss of efficacy for the ADU-S100 treatment, while the isotype group treated with a 50 µg ADU-S100 bolus consistently showed a high percentage of tumor-free survivors (two thirds, Fig. 6j). This highlights the importance of alterations in tumor infiltrating immune cells after STING agonist treatment (Fig. 3e,f,g), and the critical reshaping of the TME with profound infiltration by myeloid and NK populations, the latter being responsible for tumor rejection.

## Discussion

Here we report the immune effects and efficacy of STING activation in the brain for GBM, using a comprehensive flow cytometry panel of immune markers and intracranial gel implants for extended release of the STING agonist, to limit acute toxic effects and allow for combined therapies that overcome immunosuppressive effects over an extended period. Our data show both efficacy and feasibility of this approach. First, single IT treatment with an ADU-S100 bolus triggers a deep remodeling of the TME and rapidly drives innate infiltrates to the tumor, mainly composed of inflammatory and immature innate cells, such as macrophages, neutrophils and NK populations. NK cells have recently been highlighted as a major cytotoxic component of STING therapy (58), and although tumor regression from STING agonists has primarily been attributed to CD8+ T cells (42, 43, 72), recent data demonstrates the importance of NK cell populations in this effect, which in numerous cancer models can be CD8-independent (58, 59, 73). We here confirm the crucial participation of NK cells in the anti-tumor effects of STING agonists *in vivo*, both from the analysis of the TME in intracranial models and from the complete loss of efficacy of STING agonist treatment in NK-depleted animals. An important observation supporting the application of STING agonists in GBM comes from our use of tumor explants, which were all responsive to STING agonists as determined by inflammatory cytokine production.

The status of the STING pathway in GBM is poorly understood. STING pathway components are largely present in tumor cells and tumors in general as detected by Western blotting. Using ELISA assays we were able to show that cultured tumor cells did not respond to STING agonist treatment, with the notable exception of GL261. This lack of response in tumor cells has also been seen in other tumor types, such as melanoma for which the STING pathway is can be epigenetically silenced through cGAS or STING itself (74). Stromal cell types tested, including brain endothelial cells and pericytes were responsive to STING agonists. Our immunostaining showed that STING is present prominently in the vasculature of tumors as well as normal brain (50). Presumably this enrichment of STING provides a sensing mechanism for blood-borne pathogens in the circulation. Interestingly, we were able to detect phospho-TBK1 in tumor endothelium but not in normal brain, indicating some level of baseline pathway activation in GBM. It is unknown at present how this is stimulated, and what role it may play in GBM tumor progression, though we have previously shown that low-level activation of vascular STING by tumor-derived 2’3’-cGAMP can serve to limit T-cell infiltration in non-small cell lung cancer (50). In other brain studies, several neuropathologies have recently been linked to the activation of the STING pathway and the IFN pro-inflammatory response, such as Gaucher disease (75), Aicardi-Goutières syndrome (76) and models of prion diseases (77). Inflammatory populations like macrophages, DCs and neutrophils, which were an important part of the immune profile of the STING-agonist treated tumors in the present study, have until now been better studied in the case of brain injury and damage (57, 78, 79). STING seems to play a role in modulating immunological responses in the brain, and both neurotoxic and neuroprotective effects have been observed (57). The activation of the cGAS-STING pathway is a component of brain injury after ischemic stroke and appears deleterious after traumatic brain injury (57, 78). It is worth noting that NK cell populations here again form a central component of the tissue response after intracerebral hemorrhage and its associated inflammation (79).

Based on our work and these reports on inflammatory processes in the brain involving macrophages, DCs, neutrophils and NK populations, it seems evident that the responses are similar. It appears crucial to be able to modulate this inflammatory response and control it, notably with the use of delivery systems and combined therapies, allowing for the activation of STING and the type I IFN response to trigger sufficient but not deleterious inflammation, with the additional use of ICB or other therapies to further modulate the immune response over time. Toward that goal, we used biodegradable cross-linked gels *in vivo* and showed the feasibility of the approach. Indeed, a single treatment of GL261 and CT-2A tumors with gel-based biodegradable implants led to a cure and long-term immunity for the GL261 model and to a very significant increase in survival in the CT-2A model. The unusual CXCL10 response of GL261 cells to STING agonists, and the partial activation of the STING pathway *in vivo*, as evidenced by phosphorylation of TBK1, may contribute to the immunogenic nature of GL261 tumors in mouse, and to the high survival benefit seen in multiple preclinical immunotherapy studies in this model (18, 46, 71). Thus, intrinsic STING activation in GL261 tumors may contribute its notable immunogenicity. IFN type I generate antitumor immunity and spontaneous T cell response in both carcinogen-induced and transplantable tumor models (26, 41), and STING represents a major pathway for spontaneous antitumor immunity (25, 80, 81). It is worth noting that the GL261 tumor model was originally induced by the carcinogen methylcholanthrene in a C3H mouse, then transplanted and maintained in C57BL/6 mice (82); GL261 expresses basal major histocompatibility complex MHCI levels, but not MHCII, and carries point mutations in the K-ras and p53 genes. GL261 is considered as moderately immunogenic, while the presence of surface B7 costimulatory molecules may render them more susceptible to class I MHC-dependent cytotoxic T cells (83, 84).

The increase in median survival for the CT-2A implanted mice (18 to 29 days), although not as strong as for GL261, appears promising in light of previous reports and warrants deeper investigation of combined therapies (68). This therapeutic anti-tumor effect in CT-2A, which are non-responsive to STING agonist *in vitro*, is likely to arise from IFN production in the vasculature, with possible involvement of DCs, astrocytes and microglia (85–88). A first proof-of-concept study presented here, using a combination of ADU-S100 and anti-PD1 in the intracranial gel delivery system, shows that the combination brings more therapeutic potential than the STING therapy alone, with long term survivors in the cold and non-immunogenic CT-2A mouse model. The observation of PD-L1 upregulation in the TME post-STING agonist treatment further supports the use of ICB with anti-PD1, as does our pilot study showing long-term survival in CT-2A tumor implanted animals with anti-PD1 incorporated into the gels. The immunostimulatory effects of STING agonists in our models is also supported by transcriptomic analysis of treated GL261 tumor associated immune cells which showed a strong upregulation of IFN signaling pathways.

To summarize, our promising data confirms the importance of NK populations in the antitumor effects of STING therapies and lays the foundation for the use of STING-loaded brain implants to reshape the TME, trigger immune infiltration and serve as a support for combination therapies (ICB, cytotoxic chemotherapy) in the near future.

## Methods

### Cell culture and chemicals

GL261Luc2 murine glioma cells were purchased from PerkinElmer. CT-2A murine glioma cells were a gift from Thomas Seyfried (Boston College, Boston, Massachusetts, USA). These two murine glioma cell lines were cultured in Dulbecco’s modified Eagle’s Medium (Life technologies), containing 10% heat-inactivated fetal bovine serum (FBS) and 1% penicillin-streptomycin. HBMEC/D3 and HBVP cells were purchased from ScienCell Research Laboratories and grown according to the manufacturer’s recommendations. Primary human GSCs (G9) were obtained by dissociation of gross tumor samples and cultivated in neurosphere media (briefly: neurobasal medium containing vitamin A depleted B27, 1% Glutamax, 20 ng/mL EGF, 20 ng/mL FGF, Primocin and Plasmocin), as previously described (89). All cells were cultured at 37°C with 5% CO_2_ and mycoplasma testing was routinely done by PCR. The STING agonists ADU-S100 and 2’,3’-cGAMP were purchased from ChemieTek as the disodium salts (Cat# CT-ADUS100 and CT-CGAMP).

### Brain TMAs

Brain tumor tissue microarrays (GL2082) were purchased from US Biomax, Inc and IHC for STING and phospho-TBK1 was performed on the Leica Bond III automated staining platform as previously published (54). The antibody for phospho-TBK1 (Cell Signaling Technology #5483, clone D52C2) was run at 1:50 dilution using the Leica Biosystems Refine Detection Kit with EDTA antigen retrieval. The antibody for STING (Cell Signaling Technology #13647, clone D2P2F) was run at 1:50 dilution using the Leica Biosystems Refine Detection Kit with citrate antigen retrieval. IHC staining was quantified using the QuPath software (0.2.0-m4). The Positive Pixel Detection analysis was used with default settings for DAB staining to detect and quantify positive pixels in each of three individual, randomly selected fields per tumor, which were then averaged.

### Cytokine Analysis

CXCL10 ELISA (no. DIP100; R&D Systems) was used on media collected from cell culture according to manufacturer’s instructions. Multiplex cytokine arrays were performed as previously described (54) utilizing the bead-based immunoassay approach Bio-Plex Pro™ Human Cytokine 40-plex Assay (Cat# 171AK99MR2) on a Bio-plex 200 system (Cat# 171000201) (Bio-Rad Laboratories, Hercules, CA) and the Human Cytokine/Chemokine Magnetic Bead Panel (Cat# HCYTMAG-60K-PX30) on a Luminex MAGPIX system (Merck Millipore, Billerica, MA). Conditioned media concentrations (pg/mL) of each protein were derived from 5-parameter curve fitting models and plotted as log_2_ fold-change. Lower and upper limits of quantitation were imputed from standard curves for cytokines above or below detection. Above assay readouts are marked with asterisks.

### PBMC-mediated cytotoxicity assays

PBMCs were obtained from healthy human donors as approved by the IRB at the Brigham and Women’s Hospital (all samples were de-identified), and isolated using the Ficoll Paque Plus density gradient medium (GE Healthcare Life Sciences) following the manufacturer’s instructions. A single cell suspension of GBM cells was seeded at 2000 cells/well (G9^pCDH^) in ultra-low attachment 96-well plates (Corning) and incubated for 24 h to allow sphere formation. PBMCs were then added with the different ADU-S100 concentrations and treatment was repeated 3 days later. Cells were then incubated for another 3 days (total = 6 days of co-culture). Microscope images of the spheres were taken daily (Nikon TI, 4x magnification) and the spheres fluorescence were measured using ImageJ with a dedicated macro.

### Biodegradable cross-linked gels

The two polymeric components were prepared according to our previously published procedures (90). ADU-S100 was loaded by using an appropriate PBS solution of the drug to dissolve the norbornene-alginate component. Both components were mixed right before intracranial injection.

### *In vivo* studies

7-8 week-old female C57/BL6 mice were purchased from Envigo and a suspension of fifty thousand cells (GL261Luc2, CT-2A) in 2 μL PBS was injected intracranially to establish mouse brain tumors (2 mm right lateral, 1 mm frontal to the bregma, and 3 mm deep). Successful tumor implantation was verified by bioluminescence imaging using the Perkin-Elmer IVIS Lumina 3 system. The study end-point was considered as a weight loss of 20%, onset of neurological symptoms or signs of pain and distress. All animal experiments and procedures described in this study were approved by Brigham and Women’s Institutional Animal Care and Use Committee (IACUC).

### Isolation of murine brain-infiltrating leukocytes (BILs)

The tumor-bearing cerebral hemisphere was collected from each mouse at the indicated days after tumor implantation and therapy. Single mouse tumor cell suspensions were obtained using a mouse Tumor Disassociation Kit from Miltenyi Biotec (Cat# 130-096-730). After leukocytes extraction using density gradient medium, cell suspensions were stained and analyzed by flow cytometry.

### Mouse Tumor Flow Cytometry

Flow cytometry on mouse tumors was performed as previously described (54). Briefly, after BILs isolation, cell suspensions were subjected to flow cytometry. After live/dead staining with the Zombie NIR Fixable Viability Kit (Cat# 423106, Biolegend, San Diego, CA) per manufacturer’s instructions, single cell suspensions were stained with fluorophore-conjugated primary antibodies (see Supplementary Table S1), in PBS containing 2% FBS at 2 μg/mL. After washing, cells were resuspended in PBS containing 2% FBS and analyzed on a LSRFortessa flow cytometer (Becton Dickinson, Franklin Lakes, NJ). Levels were compared with isotype control antibodies. The data analyses were performed using the FlowJo software (TreeStar). t-SNE was achieved with the embedded FlowJo t-SNE algorithm using default parameters. FlowSOM analysis was performed with the corresponding FlowJo plugin and R. Samples of the gating strategies can be found in the Supplementary Figure S10.

### Depletion of peripheral blood NK cells

The anti-NK1.1 depleting antibody was purchased from Bio X Cell (Cat# BE0036), first injected 36 h before the intracranial treatment and biweekly after that. The mock groups used a mouse IgG2a from the same manufacturer (Bio X Cell, Cat# BE0085). All antibodies were injected intraperitoneally (250 µg in 100 µL of *InVivo*Pure pH 7.0 Dilution Buffer; Bio X Cell, Cat# IP0070). Peripheral blood was harvested from the tail vain 32 h after the first injection of the depleting antibody and was analyzed by flow cytometry to determine the NK cell contents.

### Histology

Mice were transcardially perfused with phosphate buffered saline (PBS, ThermoFisher). Brains were removed and stored 24 h in 4% PFA, then sucrose and slices were prepared using a vibratome (Campden Instruments) and immunostained for calretinin using the following solutions and protocol: carrier solution, 1% normal horse serum (NHS, Vector Laboratories) with 0.5% Triton (ThermoFisher) in phosphate buffered saline (PBS, ThermoFisher); blocking solution, 10% NHS with 0.5% Triton in PBS. After several rounds of PBS washes, slices were blocked for two hours at room temperature and incubated with primary antibody in carrier solution overnight at 4 °C. Slices were washed again in PBS and incubated for 2 h at room temperature in secondary antibody in carrier solution at a 1:1000 dilution. After four final washes, slices were mounted on slides and cover slipped using 22 × 50 mm, 0.16–0.19 mm thick cover glass (FisherScientific). Images were acquired with a LSM710 confocal microscope (Zeiss) and stitched with Zen 2.1 (black, Zeiss). Confocal images were post processed with ImageJ (Version: 2.0).

### Immunohistochemical studies and multiplexed immunofluorescence

Immunofluorescent multiplex staining was performed on the Leica Bond RX automated staining platform using the Leica Biosystems Refine Detection Kit. Antibodies were used as follows: pTBK1 (Cell Signaling Technologies, clone D52C2, Cat#5483) was run at 1:50 dilution with EDTA antigen retrieval; STING (Cell Signaling Technologies, clone D2P2F, Cat# 13647S) was run at 1:50 dilution with citrate antigen retrieval; CD31 (Cell Signaling Technology, clone D8V9E, Cat#77699) was run at 1:100 dilution with citrate antigen retrieval; IBA1 (Wako, polyclonal, Cat# 019-19741) was run at 1:500 dilution with citrate antigen retrieval. Imaging was performed on the Leica Versa 200 automated fluorescent/brightfield scanner at 20x magnification. Alexafluors 488, 555, 594 and 647 were used for each antibody (respectively).

### Immunoblotting

Proteins were isolated from cell lines and content measured by BCA (Pierce Biotechnology). Protein extracts were subjected to polyacrylamide gel electrophoresis using either Criterion or Mini-Protean TGX pre-cast gels, transferred to nitrocellulose (Millipore) membranes, and immunoblotted using antibodies (Cell Signaling Technologies, Danvers, MA) that specifically recognize STING (clone D2P2F, Cat# 13647S), cGAS (clone D1D3G, Cat# 15102), TBK1 (clone E8I3G, Cat#38066), human or mouse anti-GAPDH (ab9484-200, Abcam, Cambridge, MA, USA) and peroxidase-conjugated secondary antibodies (Jackson Laboratories, Bar Harbor, ME, USA). 5% BSA blocking buffer was used to dilute primary and secondary antibodies. Imaging of blots and quantitation of bands was performed using the Biorad Gel Doc XR Imaging System.

### Imaging methods

MRI data were acquired using either a BioSpec 3T or a Bruker 7 Tesla. Animals were kept under isoflurane narcosis throughout the scan. Respiration and heart rate were monitored. T2-weighted images were acquired using the RARE pulse sequence with the following settings: TE (echo time): 47.73ms, TR (repetition time): 4993.715ms, Rare Factor: 8, Averages: 3. Slice thickness: 0.5mm, slicer orientation: axial. Field of View: 20mm * 20mm, Resolution: 0.078mm * 0.078mm.

### Patient samples

The brain tumor samples were collected under the institutional banking IRB approved protocol 10-417. The samples were distributed under tissue sub usage protocol approval. All patients undergoing a brain tumor surgery at the Brigham are open to this banking protocol at the time of surgery. The IRB is approved by the DF/HCC IRB and signed consent was obtained from all patients. Freshly isolated tumor tissue was harvested and immediately processed within a few hours of surgery.

### RNAseq from BILs

Total RNA was isolated directly from triplicate samples (BILs three days after ADU-S100) with the Qiagen (Hilden, Germany) RNeasy isolation kit (Cat# 8028) using on-column DNAse digestion. RNA libraries were prepared from 250 ng total RNA using the Illumina Exome Capture kit per manufacturer’s instructions. RNA-sequencing (RNA-seq) was performed per the standard protocols at the Dana-Farber Molecular Biology Core Facilities with Illumina NextSeq 500 instrument. Data quality controls and replicate correlation were evaluated using VIPER software package.(91) Briefly, libraries were prepared using SMARTer Stranded Total RNAseq v2 Pico Input Mammalian sample preparation kits from 500 pg of purified total RNA according to the manufacturer’s protocol. The finished dsDNA libraries were quantified by Qubit fluorometer and Agilent TapeStation 4200. Uniquely dual indexed libraries were pooled in an equimolar ratio and shallowly sequenced on an Illumina MiSeq to further evaluate library quality and pool balance. The final pool was sequenced on an Illumina NovaSeq 6000 targeting 80 million 100 bp read pairs per library at the Dana-Farber Cancer Institute Molecular Biology Core Facilities.

### Bulk RNA-Seq Analysis

Sequencing reads were aligned using STAR (92) with an average of 3E7 uniquely mapped reads per sample and a mismatch rate per base of less than 1%. Aligned reads were first filtered to remove genes with less than 10 counts across all samples. Differential expression analysis was performed using DESeq2 in R (93), with FDR adjusted p-value threshold of <=0.1. A volcano plot showing log_2_ fold change and -log_10_ (adjusted p value) was then generated from the differential expression analysis using ggplot2. Gene enrichment analysis was performed using Enrichr (94) to investigate enriched pathways. The -log_10_ (adjusted p value) of the most significant Gene Ontology (Biological Process) pathways for each condition were then plotted as bar graphs, alongside the corresponding values for the other condition for comparison. To further compare IFN gene signatures between ADU treated samples and PBS controls, gene sets known to be responsive to IFNα, IFNβ and IFNγ treatment were curated from published literature (63). The distributions of gene expression for the various IFN gene signatures per treatment group were estimated by first applying a regularized log transformation the filtered gene counts, followed by z-score normalization by gene. Violin plots were then used to visualize the distribution of the normalized gene expression values by condition. Statistical comparisons were performed using Wilcox test, with FDR for p-value adjustment.

### Statistical analyses

All graphs depict mean ± s.e.m unless otherwise indicated. Tests for differences between two groups were performed using unpaired two-tailed Student’s t-test. Multiple comparisons used one-way or two-way ANOVA, as specified in the figure legends. Log-rank test was utilized for patient and mouse survival analyses. GraphPad Prism 9 was used for statistical analysis of experiments, data processing and presentation. Sample sizes for *in vivo* studies were determined empirically based on results from prior publications.

## Author contributions

Conceptualization, G.B., E.H.K. and S.E.L.; Experimental support, M.O.N, S.H., P.H.L., J.L.L., A.S., N.D.; RNAseq analysis A.K.-B.; Original draft, G.B. and S.E.L.; Revisions, all authors; Supervision and funding, S.E.L.

## Acknowledgments

We thank Dana-Farber/Harvard Cancer Center in Boston, MA, for the use of the Specialized Histopathology Core, which provided histology and immunohistochemistry service. Dana-Farber/Harvard Cancer Center is supported in part by an NCI Cancer Center Support Grant # NIH 5 P30 CA06516. Additional funding by Expect Miracles Foundation and the Robert A. and Renée E. Belfer Family Foundation. The authors thank Tomer Filkenberg for his help on IF staining and Brandon Piel for GBM explants.

