## Supplementary Data for "STING activation promotes robust immune response and NK cell-mediated tumor regression in glioblastoma models"

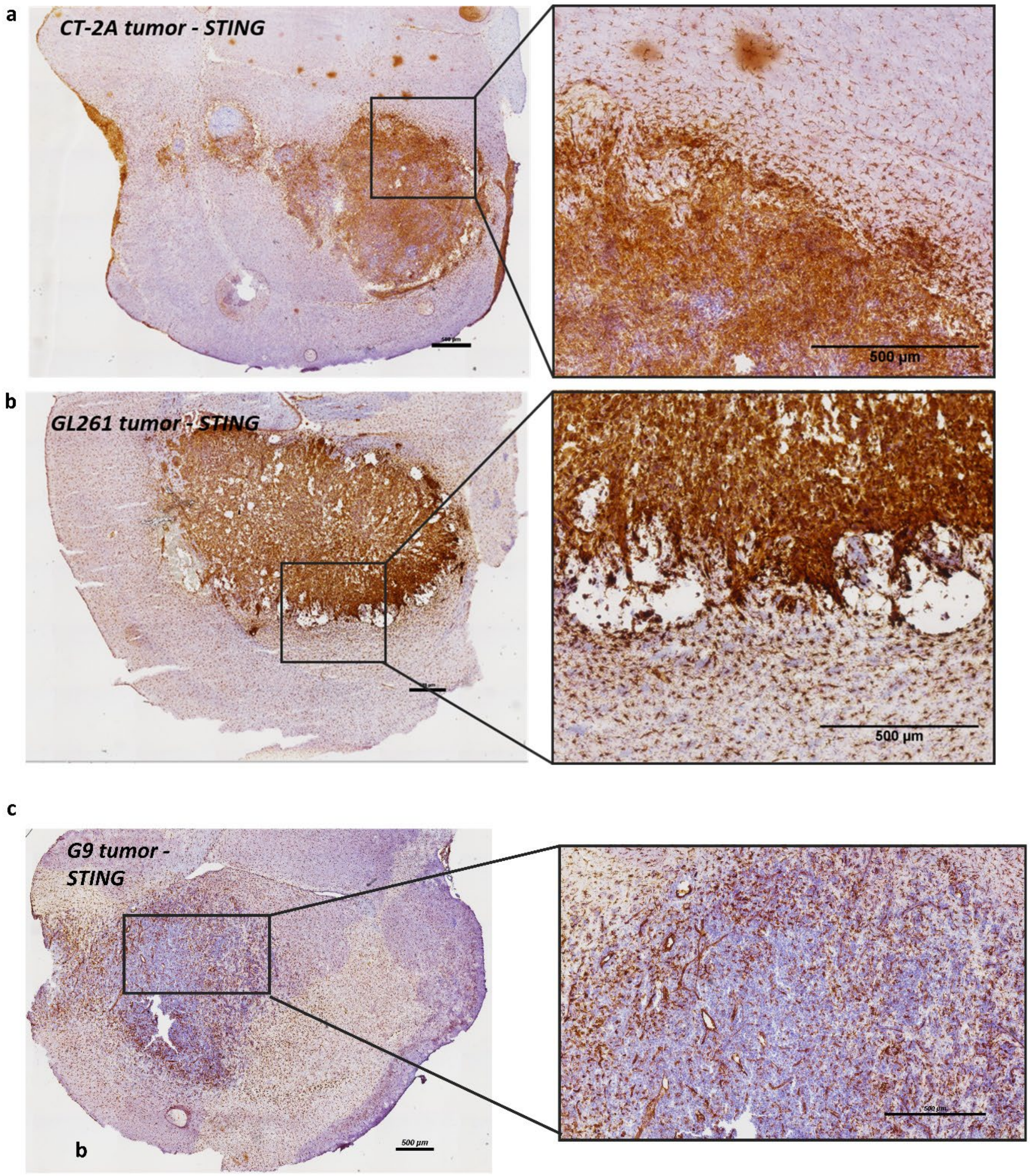


**Figure S1. STING immunostaining on mouse tumor models. a, b,** STING IHC staining of established murine CT-2A and GL261 tumors, showing STING is widely expressed in the tumor and expressed in a subset of cell in the healthy tissue. **c,** STING IHC staining for a G9 xenograft tumor; the bulk of the tumor does not express STING to visible levels.

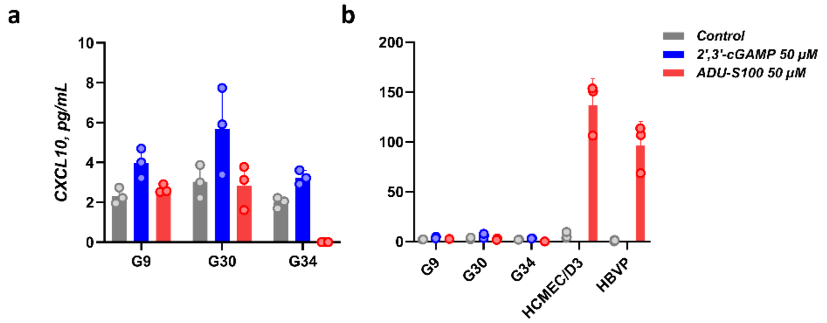


**Figure S2. CXCL10 production following STING activation by 2',3'-cGAMP and ADU-S100.** **a**, Levels of CXCL10 as measured by ELISA 24 h after STING agonist treatment of the indicated human GBM cell neurosphere lines. **b**, The same graph with the addition of responsive human brain endothelial HCMEC/D3 and brain pericyte HBVP cells to allow comparison.

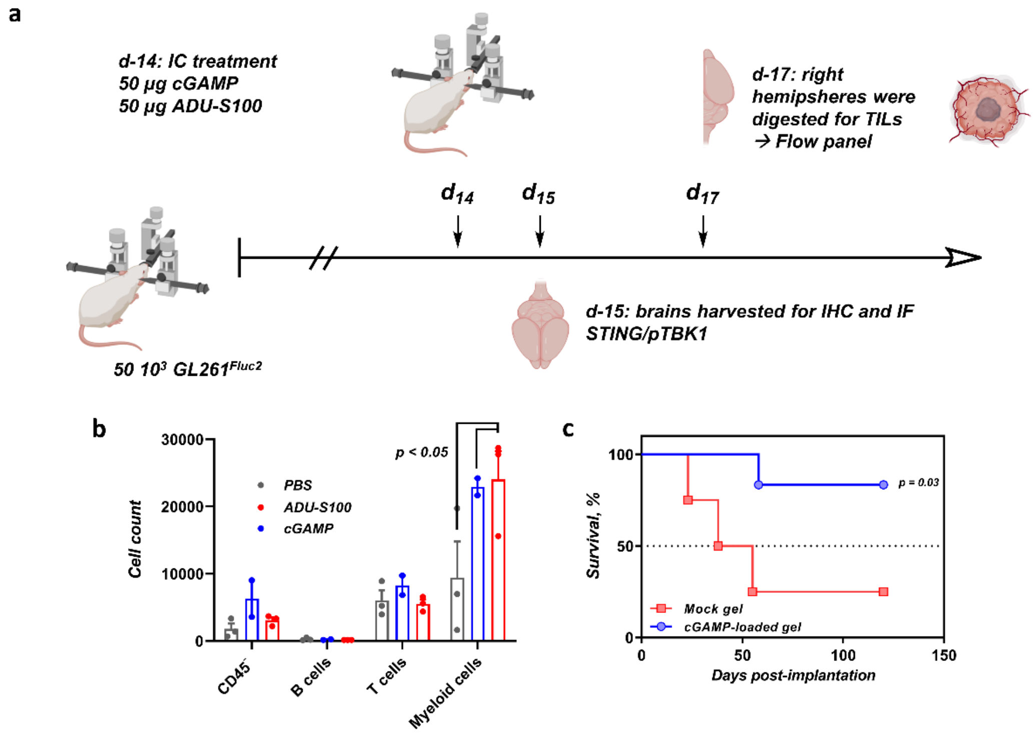


**Figure S3. Pilot *in vivo* STING experiment. a**, Timeline of the experiment. **b**, Flow cytometry analysis of the BILs 3 days after treatment with a bolus of either cGAMP or ADU-S100 in PBS (50 µg). **c**, Kaplan-Meier survival analysis from a cohort implanted with cGAMP-loaded hydrogels (100 µg).

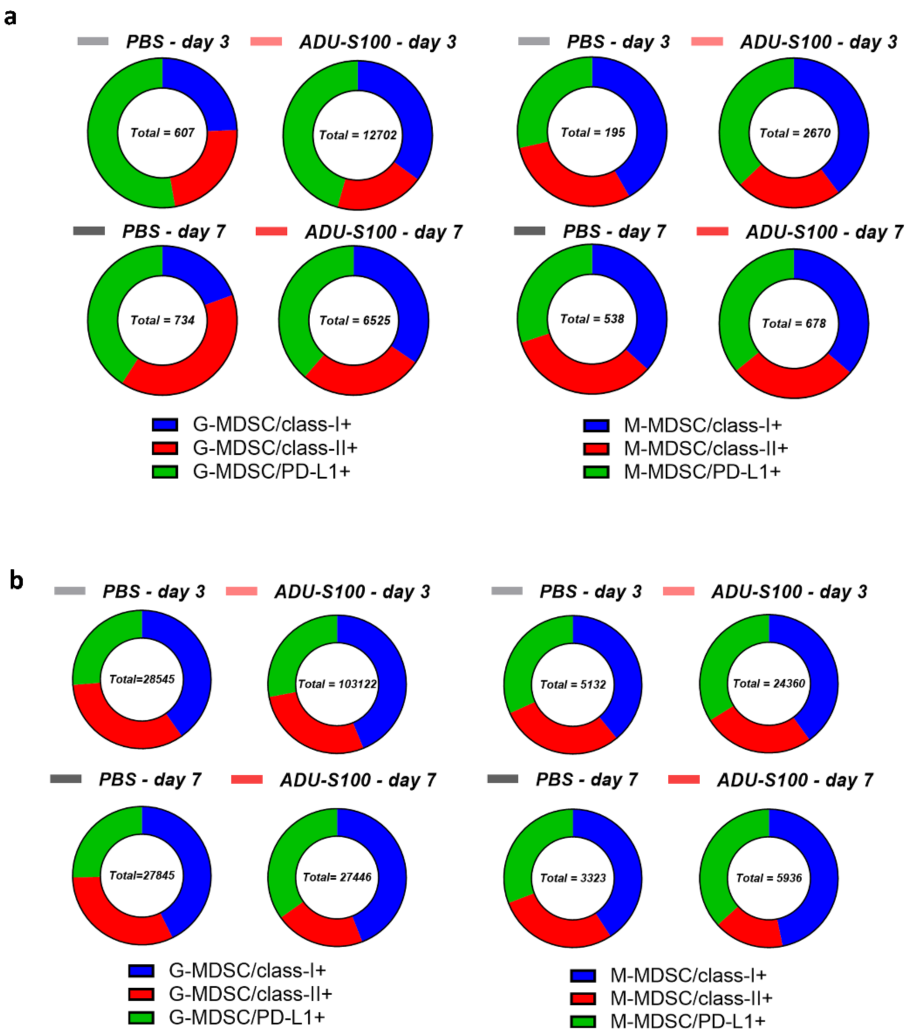


**Figure S4. Flow cytometry analysis of brain MDSC populations following STING activation by ADU-S100. a, b,** Expression levels of MHC class I/II and PD-L1 on G-MDSC and M-MDSC from BILs extracted from GL261 and CT-2A established tumors in controls and ADU-S100 treated conditions (respectively a and b).

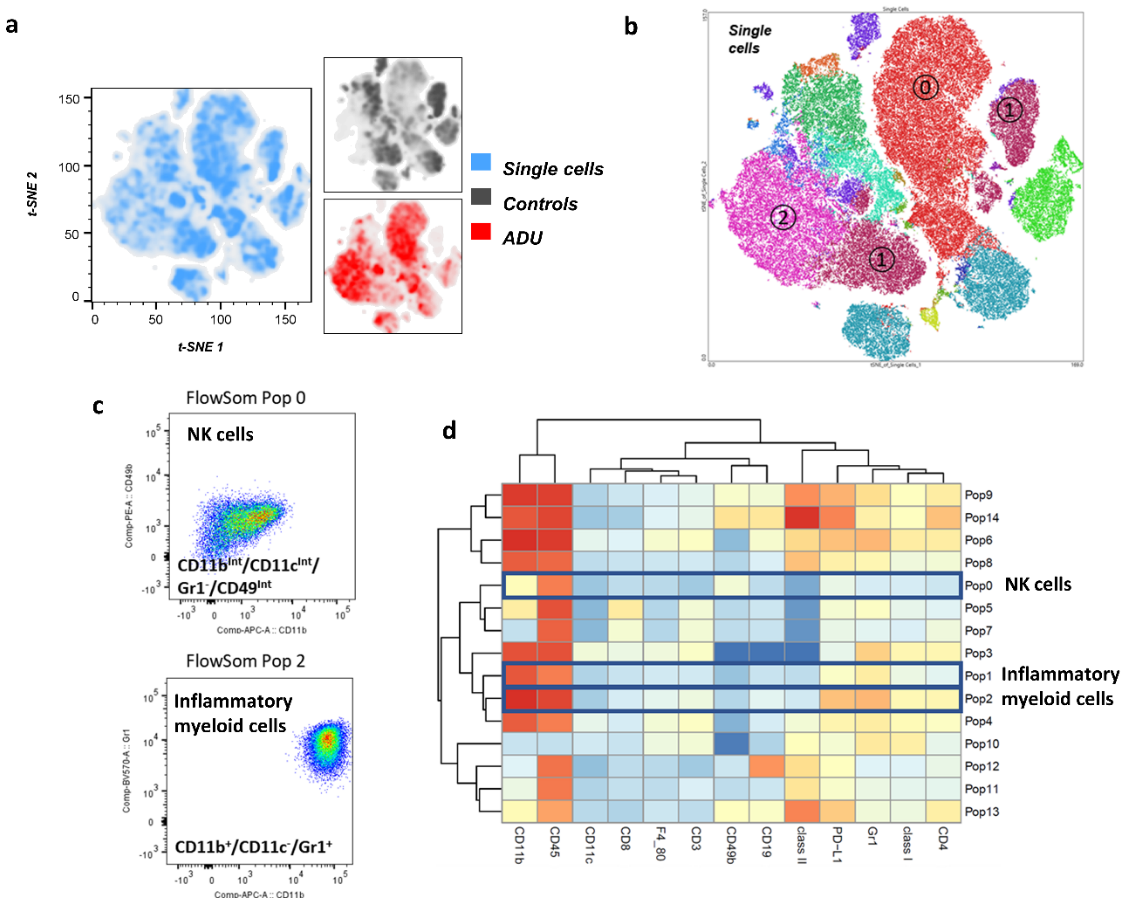


**Figure S5. Assessment of CT-2A tumor immune infiltrates: figures at day 7 after STING agonist treatment.** **a**, 2D t-SNE plots at day 7 post-treatment of established CT-2A tumors with ADU-S100 treated mice (50 µg, bolus) in red and controls in dark grey. **f**, t-SNE map for treated mice at day 7 colored by the FlowSOM populations; with relevant cell types highlighted. **c**, highly upregulated populations, comprising NK and inflammatory cells. **d,** Heatmap and hierarchical clustering of the FlowSOM populations at day 7.

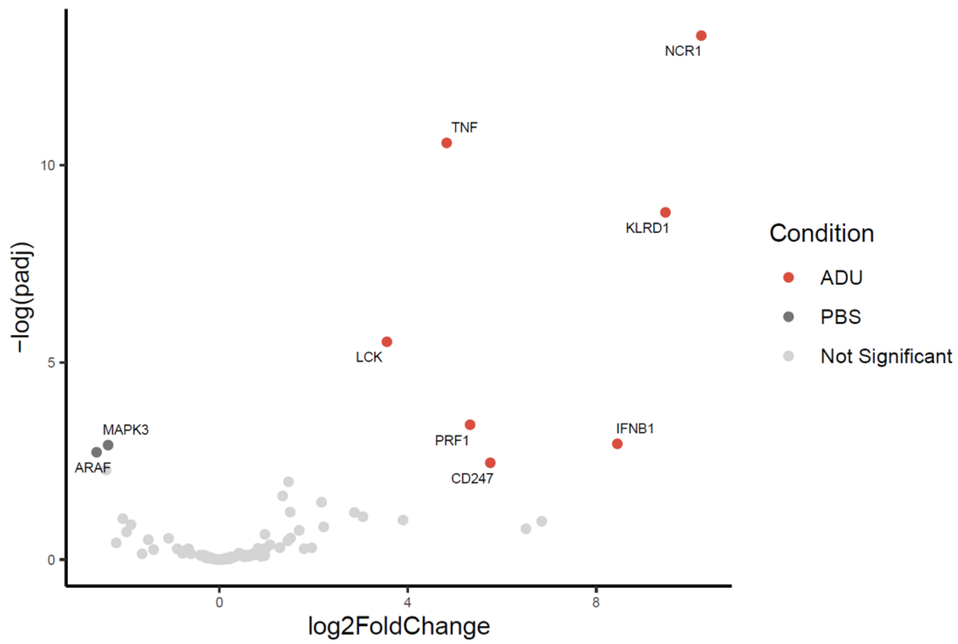


**Figure S6. Expression of NK cell associated genes in response to STING agonists.** Volcano plot for RNA sequencing data from GL261 BILs, ADU-S100 vs PBS showing differentially expressed genes for the NK mediated cytotoxicity KEGG gene set. (FDR adjusted *p* threshold ≤ 0.1.).

**Reference:**

**https://www.gsea-msigdb.org/gsea/msigdb/cards/KEGG_NATURAL_KILLER_CELL_MEDIATED_CYTOTOXICITY**

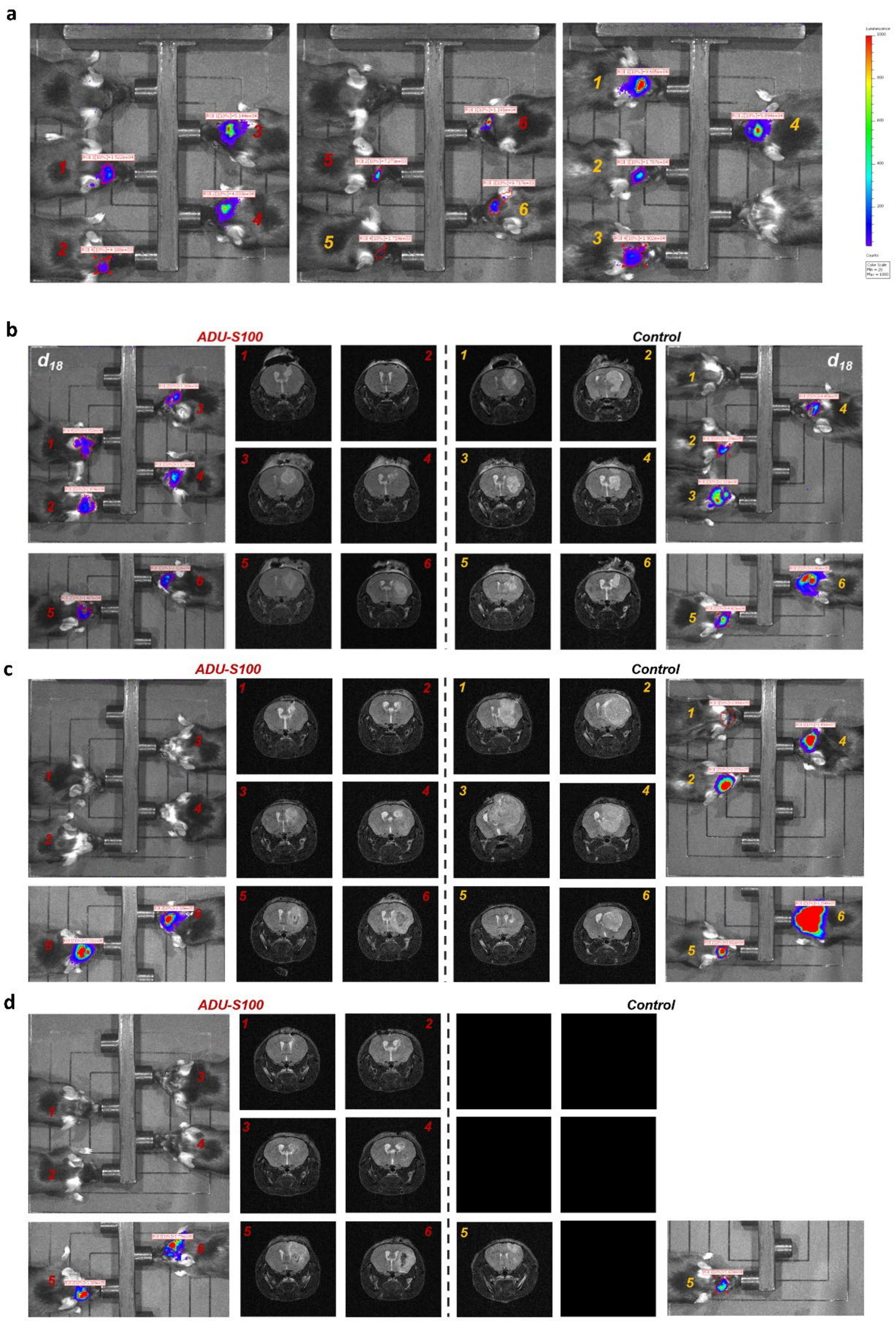


**Figure S7. Therapeutic STING implants: GL261 survival. a,** IVIS picture of both groups on treatment day. Mice without a clear IVIS signal are discarded from the study. **b, c and d,** IVIS and MRI of both groups at different timepoints.

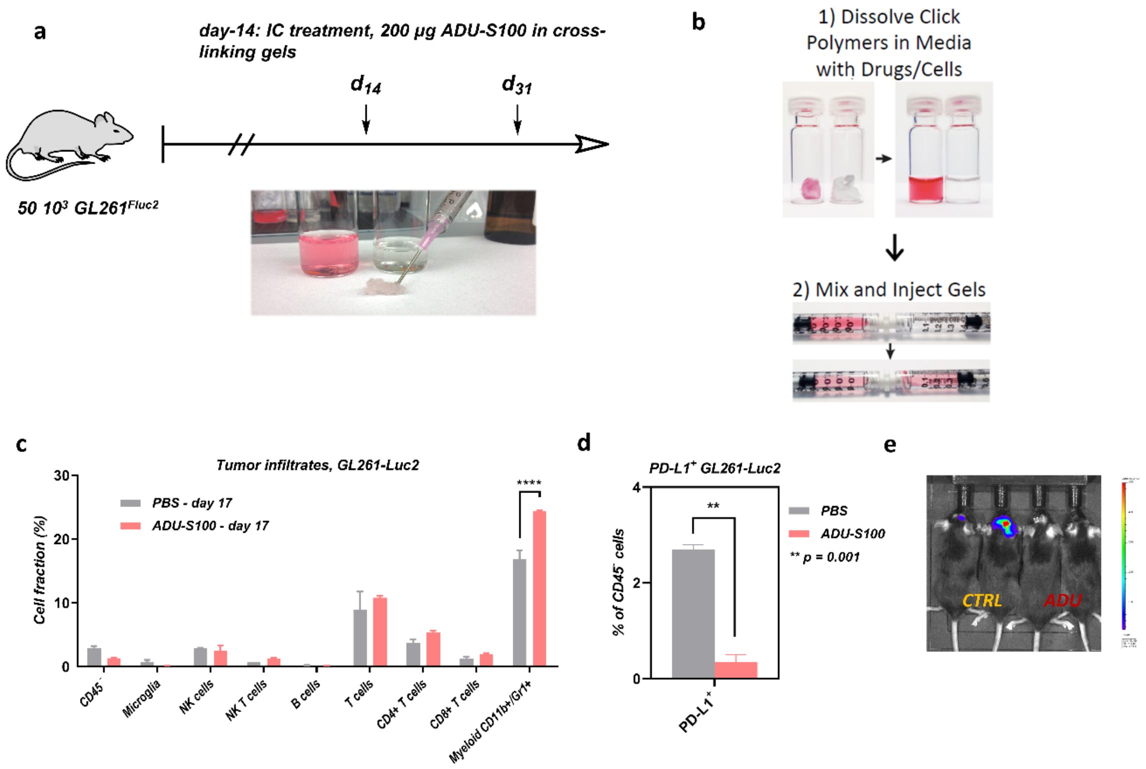


**Figure S8. Long-term effect of therapeutic STING implants on the GL261 model. a,** Timeline of the experiment. **b,** Graphical summary of the gel preparation. c, BIL flow panel 17 days after therapeutic gel implantation. **d,** PD-L1 expression on CD45⁻ cells at the same timepoint. **e,** IVIS of the mice before sacrifice.

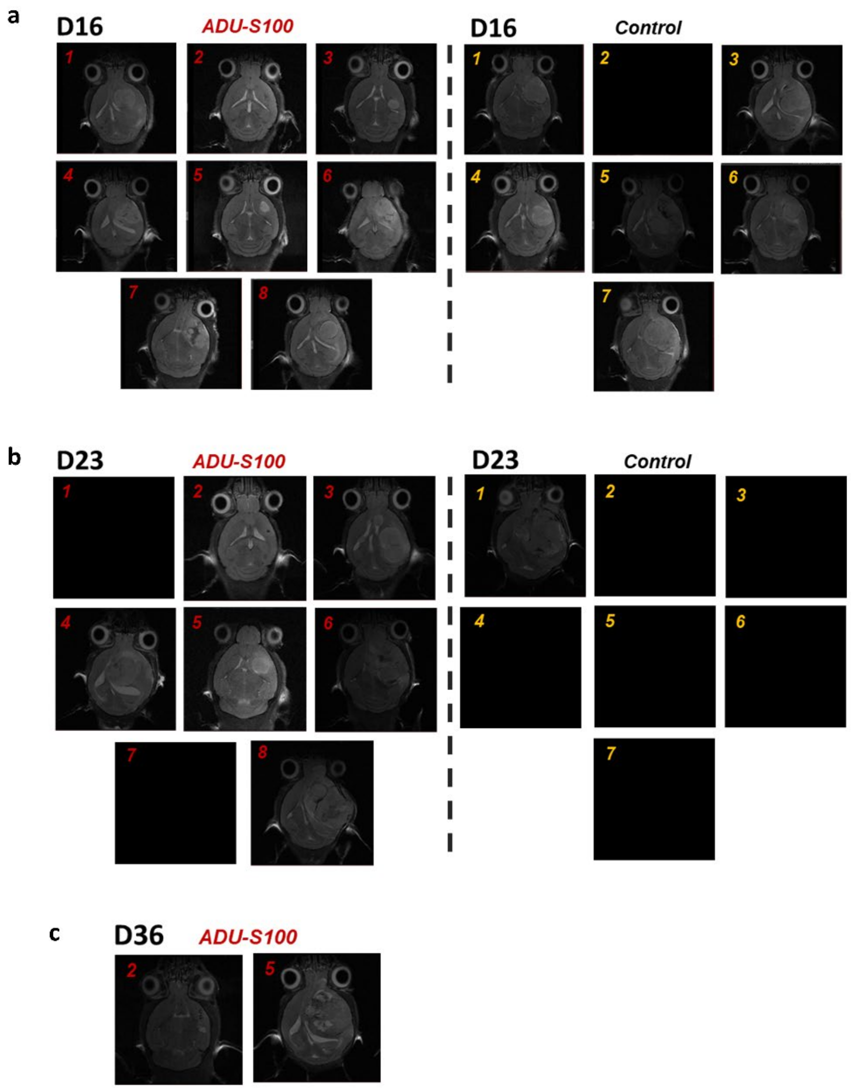


**Figure S9. Therapeutic STING implants: CT-2A survival. a, b and c,** IVIS and MRI imaging of both groups at different timepoints as shown.

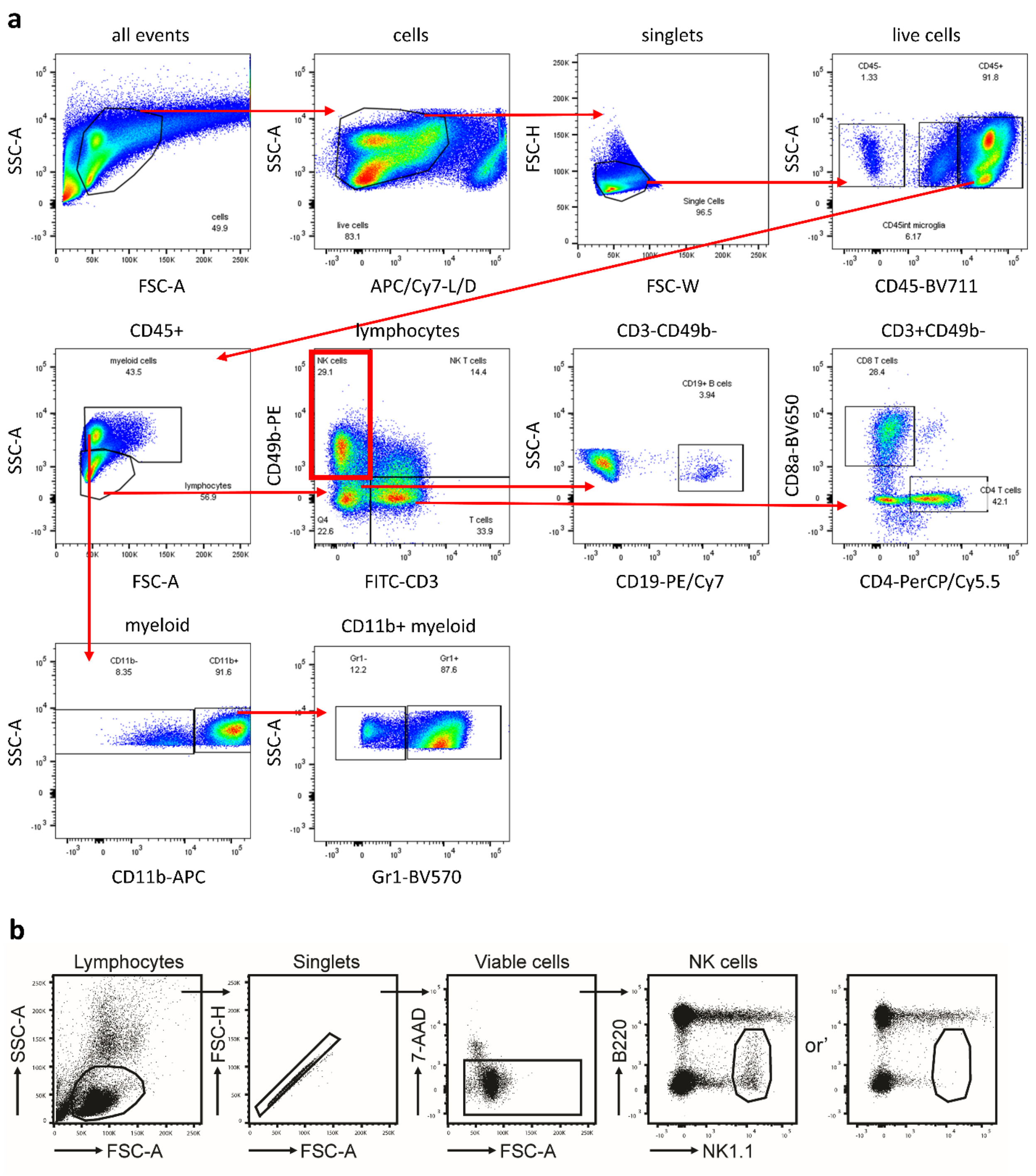


**Figure S10. a,** Gating strategies employed for the analysis of the TME and **b**, for the quantification of NK cells in peripheral blood.

**Table 1.** Mouse Flow Panel (all antibodies purchased from Biolegend, San Diego, CA).

| | |
|---|---|
| CD3 FITC 100306 | F4/80 BV421 123132 |
| CD4 PerCP/Cy5 100434 | H2KB BV510 116523 |
| CD19 PE/Cy7 115520 | GR-1 BV570 108431 |
| CD49b PE 108908 | CD8a BV650 100742 |
| PD-L1 PE Dazzle 594 124323 | CD45 BV711 103147 |
| CD11b APC 101212 | CD11c BV785 117335 |
| MHC II AF700 107622 | |
